# Paleoecological implications of Lower-Middle Triassic stromatolites and microbe-metazoan build-ups in the Germanic Basin: Insights into the aftermath of the Permian – Triassic crisis

**DOI:** 10.1101/2021.07.12.452070

**Authors:** Yu Pei, Hans Hagdorn, Thomas Voigt, Jan-Peter Duda, Joachim Reitner

**Affiliations:** Department of Geobiology, Geoscience Center, Georg-August-Universität Göttingen, Göttingen 37077, Lower Saxony, Germany; Muschelkalkmuseum, Ingelfingen 74653, Baden-Württemberg, Germany; Institute of Geosciences, Friedrich-Schiller-Universität Jena, Jena 07749, Thuringia, Germany; Sedimentology & Organic Geochemistry Group, Department of Geosciences, Eberhard-Karls-Universität Tübingen, Tübingen 72076, Baden-Württemberg, Germany; ‘Origin of Life’ Group, Göttingen Academy of Sciences and Humanities, Göttingen 37073, Lower Saxony, Germany

**Keywords:** Upper Buntsandstein, Middle Muschelkalk, Non-spicular (“keratose”) demosponges, End-Permian mass extinction, Microbialite, Ecological niche, Hypersalinity, Paleoenvironment, Geobiology

## Abstract

The aftermath of the Permian – Triassic crisis is characterized by ubiquitous occurrences of microbial sediments around the world. For instance, Triassic deposits of the Germanic Basin have shown to provide a rich record of stromatolites as well as of microbe-metazoan build-ups with non-spicular demosponges. Despite their paleoecological significance, however, all of these microbialites have only rarely been studied. This study aims to fill this gap by examining and comparing microbialites from the Upper Buntsandstein (Olenekian, Early Triassic) and the lower Middle Muschelkalk (Anisian, Middle Triassic). By combining analytical petrography (optical microscopy, micro X-ray fluorescence, Raman spectroscopy) and geochemistry (δ^13^C_carb_, δ^18^O_carb_), we show that all studied microbialites formed in hypersaline lagoons to sabkha environments. Olenekian deposits in Jena and surroundings and Anisian strata at Werbach contain stromatolites. Anisian successions at Hardheim, in contrast, host microbe-metazoan build-ups. Thus, the key-difference is the absence or presence of non-spicular demosponges in microbialites. After the Permian – Triassic crisis, the widespread microbialites (e.g., stromatolites/microbe-metazoan build-ups) possibly resulted from suppressed ecological competition and occupied the vacant ecological niche. It seems plausible that microbes and non-spicular demosponges had a mutualistic relationship and it is tempting to speculate that the investigated microbial-metazoan build-ups reflect an ancient evolutionary and ecologic association. Furthermore, both microbes and non-spicular demosponges may benefit from elevated salinities. Perhaps it was minor differences in salinities that controlled whether or not non-spicular demosponges could develop.

## 1 Introduction

Microbialites represent benthic microbial communities (i.e., biofilms or microbial mats) fossilized through trapping and binding of detrital sediment and/or localized mineral precipitation (Burne and Moore 1987; Arp et al. 2001; Riding and Liang 2005; Riding 2006). Dating as far back as ca. 3.4 billion years ago (e.g., Duda et al. 2016), they are abundant during most of the Precambrian, but show a marked decline during the Phanerozoic (Arp et al. 2001; Riding 2006). However, microbialites show marked reoccurrences at certain times in the Phanerozoic, as for instance in the aftermath of the Permian – Triassic crisis (Woods 2014; Wu et al. 2017; Heindel et al. 2018; Chen et al. 2019; Pei et al. 2019). The temporary proliferation of microbial mats in this critical interval likely resulted from a suppressed ecological competition and/or seawater chemistry at that time (Arp et al. 2001; Riding and Liang 2005; Foster et al. 2020).

Depending on their characteristics, microbialites can be classified into different types (e.g., stromatolites, thrombolites, dendrolites and leiolites: Kalkowsky 1908; Aitken 1966; Burne and Moore 1987; Riding 1991; Braga et al. 1995). Among these, stromatolite derives from the term ‘Stromatolith’, which was coined by Kalkowsky (1908) to describe layered carbonate structures in the Lower Triassic Buntsandstein of central Germany. Importantly, this term does not consider the nature of biofilms or microbial mats. In fact, the microbial communities involved in stromatolite formation might range from phototrophs (e.g., cyanobacteria) (e.g., Walter 1972; Dravis 1983; Arp et al. 1999; Bosak et al. 2012) to light-independent chemolithotrophic or heterotrophic microbes (e.g., Playford et al. 1976; Böhm and Brachert 1993; Heim et al. 2015, 2017; Mißbach et al. 2021). Particularly noteworthy are stromatolites formed by syntrophic consortia of anaerobic methane-oxidizing archaea and sulfate reducing bacteria (Reitner et al. 2005a, b; Arp et al. 2008).

Recently, a new type of microbialite formed by microbes and metazoans (mainly non-spicular demosponges) was established (Luo and Reitner 2014, 2016) and is called microbe-metazoan build-ups (Pei et al. 2021). Previously, ‘reticular fabrics’ were described by Szulc (1997) attributed to sponges in microbialites from the Middle Muschelkalk *Diplopora* Dolomite of South Poland. Unfortunately, the build-ups are easily overlooked in geological records. This is because the involved demosponges have a relatively low fossilization potential due to the absence of sponge spicules and possess hardly identifiable morphological characteristics. However, pioneering work established robust criteria for the identification of non-spicular demosponges in ancient records and demonstrated the presence of these organisms in various Phanerozoic microbe-metazoan build-ups (Luo and Reitner 2014, 2016). Afterwards, much attention and interests have been aroused (Friesenbichler et al. 2018; Heindel et al. 2018; Lee and Riding 2021; Baud et al. 2021; Pei et al. 2021).

In the Lower and Middle Triassic, the Germanic Basin recorded at least five intervals of major microbialite development, namely (i) in the Lower Buntsandstein (Kalkowsky 1908; Paul et al. 2011), (ii) in the Upper Buntsandstein (Naumann 1928), (iii) in the Karlstadt Formation of the lower Middle Muschelkalk (Genser 1930, Schwarz 1970, Vossmerbäumer 1971, Hagdorn and Simon 1993), (iv) in the Diemel Formation of the upper Middle Muschelkalk (Pei et al. 2021), and (v) in the Lower Keuper (Bachmann and Gwinner 1971; Bachmann 2002; Luo and Reitner 2016). Despite their significance for understanding the aftermath of the crisis, these microbialites have only rarely been studied. We aim to fill this gap by examining and comparing microbialites from the Upper Buntsandstein (Olenekian, Early Triassic) and the lower Middle Muschelkalk (Anisian, Middle Triassic). Based on our findings, we will discuss paleoecological implications of these records.

## 2 Geological setting

The Permian–Triassic Germanic Basin stretched across large areas of today’s Western and Central Europe. During these times, the basin was situated on the edge of the subtropical western Tethys Ocean (Meliata Ocean) (Scotese and McKerrow 1990; Ziegler 1990; Stampfli 2000) (Fig. 1). In the Olenekian (Early Triassic), it was connected to the Tethys Ocean systems through the East Carpathian Gate (ECG). During the Anisian (Middle Triassic), it was additionally linked by and the Silesian – Moravian Gate (SMG) and the Western Gate (WG) (Ziegler 1990; Götz and Feist-Burkhardt 2012; Hagdorn 2020) (Fig. 1). Paleogeographically, three of the studied sections (Jena and surroundings, Werbach and Hardheim) were located between the Rhenish Massif (RM) and the Vindelician – Bohemian – Massif (VBM) (Fig. 1).

**Fig. 1.**
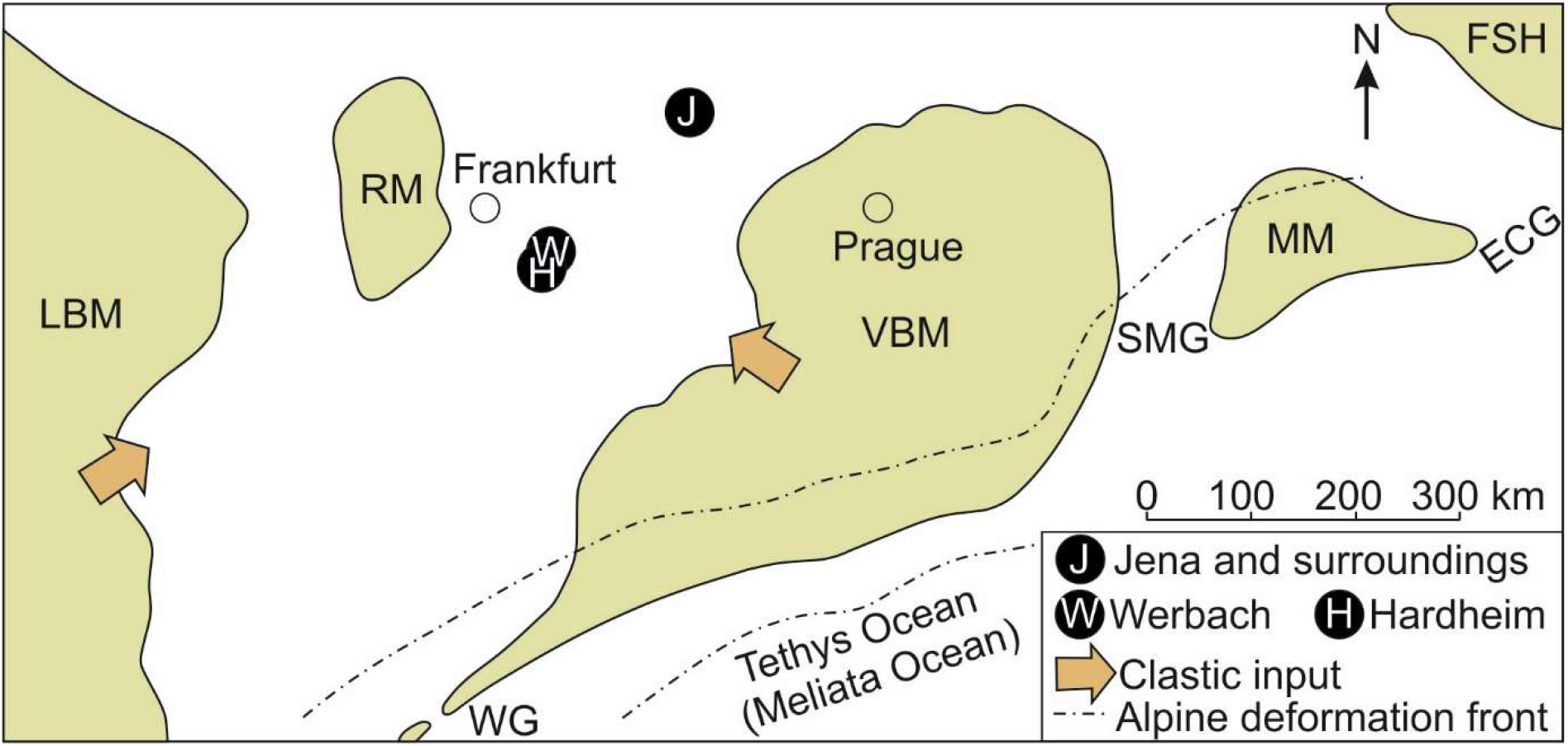
Paleogeographic map of the Germanic Basin during the Triassic (after Scotese and McKerrow 1990; Ziegler 1990; Stampfli 2000; Götz and Feist-Burkhardt 2012; Hagdorn 2020). During the Olenekian (Early Triassic), the Germanic basin was connected to the Tethys (Meliata Ocean) through the East Carpathian Gate (ECG). During the Anisian (Middle Triassic), the Germanic Basin and the Tethys Ocean systems were additionally linked through the Silesian – Moravian Gate (SMG) and the Western Gate (WG). Three of the studied sections (Jena and surroundings, Werbach and Hardheim) were located between the Rhenish Massif (RM) and the Vindelician – Bohemian – Massif (VBM). LBM: London – Brabant – Massif, MM: Malopolska Massif, FSH: Fenno – Scandian High

Sedimentary rocks in Jena and surroundings (Thuringia, central Germany) belong to the Lower Röt Formation (Upper Buntsandstein Subgroup, Olenekian, Early Triassic). The rocks are characterized by evaporites and marls to sandstones that are intercalated with dolomites and bioclastic limestones (Fig. 2). The *Tenuis*-bank at the base of the section contains abundant specimens of the ammonoid *Beneckeia tenuis*, the earliest Triassic ammonoid in the Germanic Basin, and is followed by stromatolites (Fig. 2). The researched stromatolites at Werbach quarry (Baden-Württemberg, Southwest Germany) belong to the Geislingen Bed, a supraregional marker horizon (Simon et al. 2020), in the lower part of the Karlstadt Formation of the Middle Muschelkalk Subgroup (Anisian, Middle Triassic) (Fig. 3). From bottom to top, the section can be subdivided into the Karlstadt, the Heilbronn and the Diemel formations. The Karlstadt Formation is composed of dolomites, dolomitic marls, and dolomitic limestones. Locally, it contains fossils of hypersaline organisms and stromatolites. The salinar Heilbronn Formation is almost devoid of any fossils. The Diemel Formation consists of dolomitic limestones and locally contains euryhaline faunas (Simon et al. 2020). The microbe-metazoan build-ups exposed at the abandoned quarry near Hardheim (Baden-Württemberg, Southwest Germany) occur on top of the Geislingen Bed. The build-ups are stratigraphically correlated with the stromatolites at Werbach section.

**Fig. 2.**
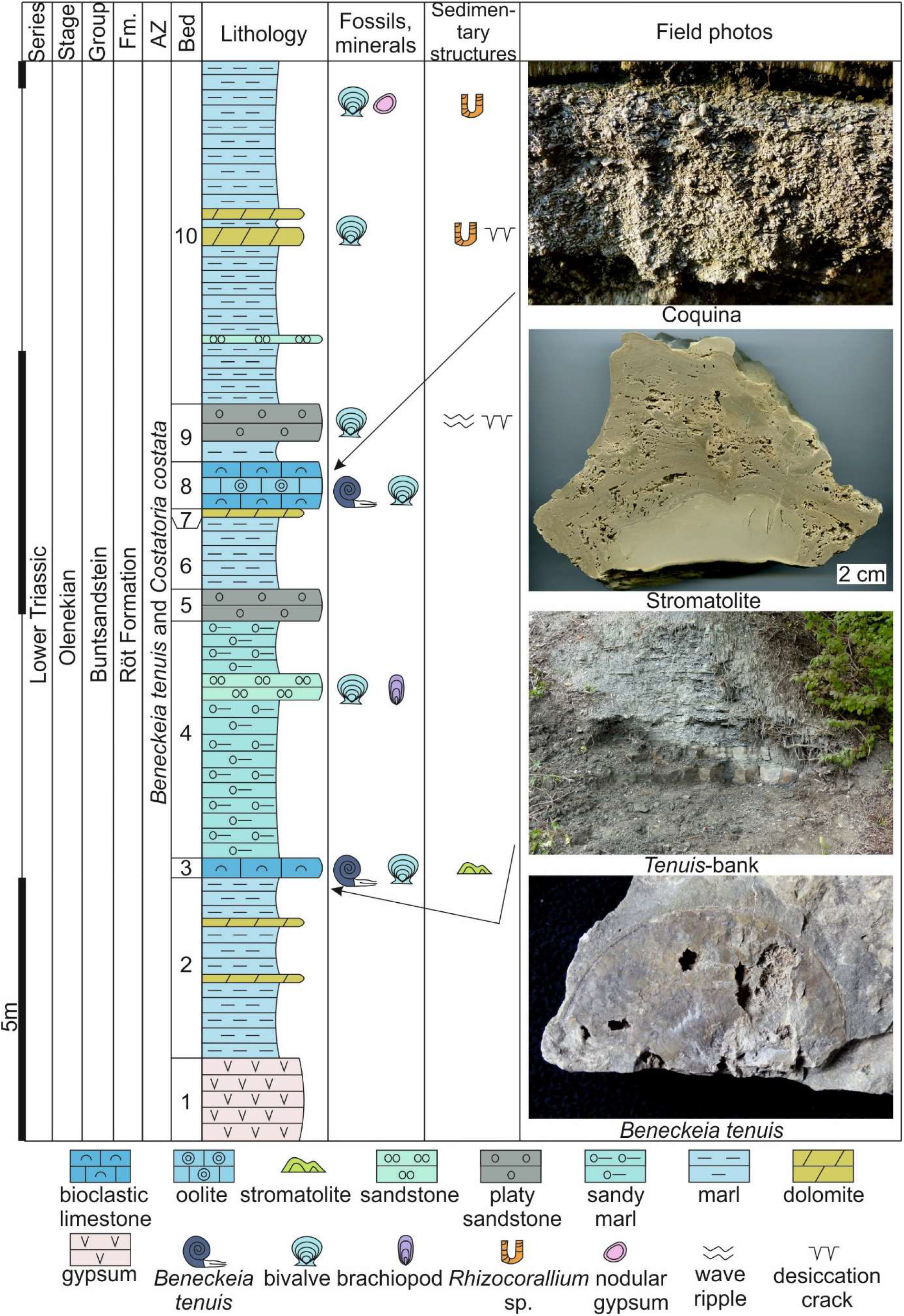
The Jena and surroundings section (Upper Buntsandstein, Olenekian, Early Triassic, Germanic Basin), showing stratigraphical, sedimentological and paleontological features

**Fig. 3.**
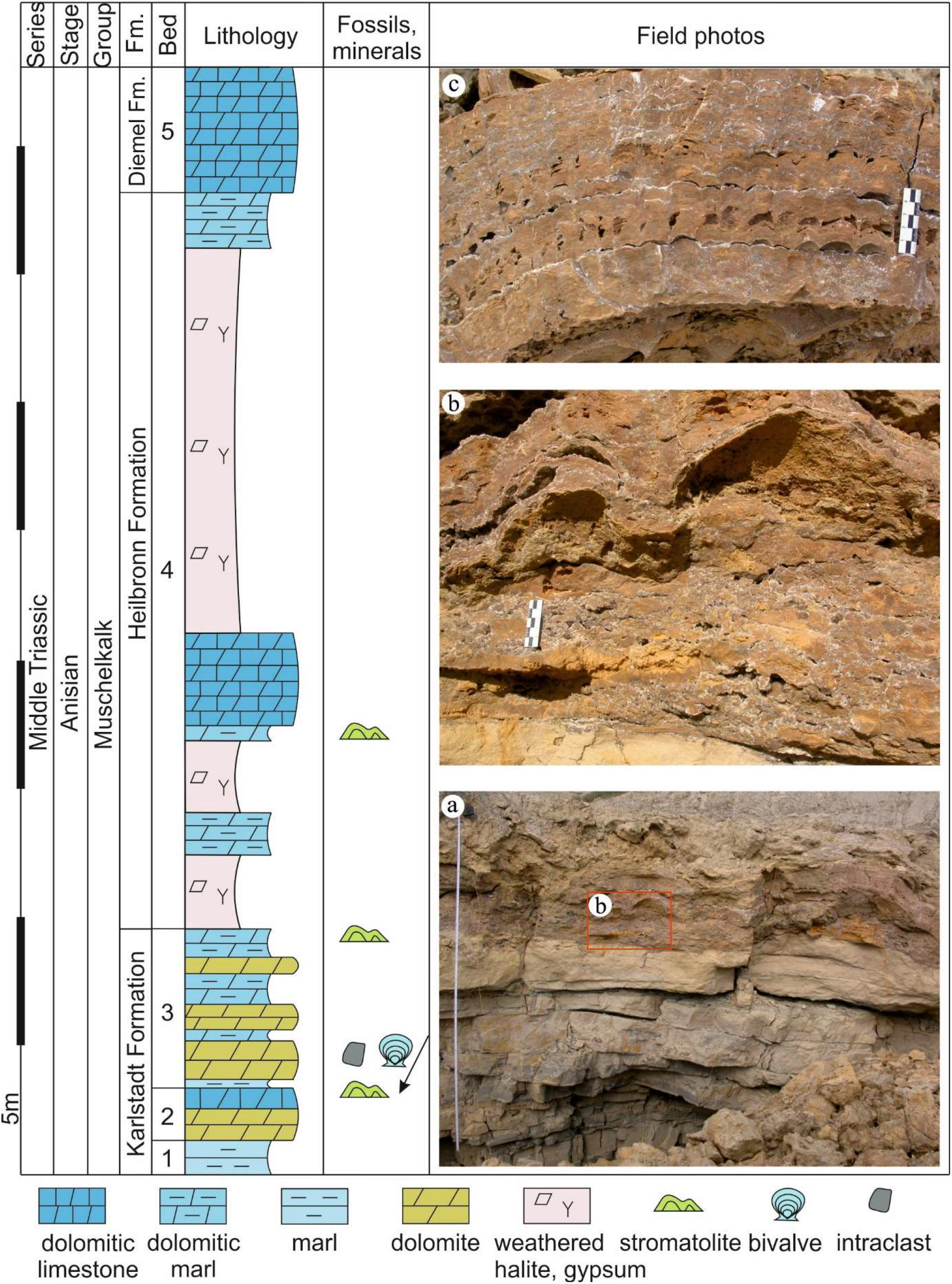
The Werbach section (lower Middle Muschelkalk, Anisian, Middle Triassic, Germanic Basin), showing stratigraphical, sedimentological and paleontological features

## 3 Materials and methods

### 3.1 Fieldwork and petrography

Sections in Jena and surroundings and at Werbach were examined in the field and fresh samples were taken (stromatolites from Jena and surroundings, stromatolites and associated facies from Werbach, and microbe-metazoan build-ups from Hardheim). Petrographic thin sections were prepared and analyzed using a Zeiss SteREO Discovery.V12 stereomicroscope coupled to an AxioCamMRc camera. The samples were then further studied by means of analytical imaging techniques and stable isotope analyses (see below).

### 3.2 Analytical imaging techniques

Micro-X-ray fluorescence (μ-XRF) was applied to obtain element distribution images of the sampled stromatolites and microbe-metazoan build-ups. The analyses were conducted with a Bruker M4 Tornado instrument equipped with an XFlash 430 Silicon Drift Detector. Measurements (spatial resolution = 25–50 µm, pixel time = 8–25 ms) were performed at 50 kV and 400 µA with a chamber pressure of 20 mbar.

Raman spectroscopy analyses included point measurements (single spectra) and mapping (spectral images). For these analyses, a WITec alpha300R fiber-coupled ultra-high throughput spectrometer was used. Before analysis, the system was calibrated using an integrated light source. The experimental setup included a laser with an excitation wavelength of 532 nm, an automatically controlled laser power of 20 mW, a 100× long working distance objective with a numerical aperture of 0.75, and a 300 g mm^−1^ grating. The spectrometer was centered at 2220 cm^−1^, covering a spectral range from 68 cm^−1^ to 3914 cm^−1^. This setup has a spectral resolution of 2.2 cm^−1^. For single spectra, each spectrum was collected by two accumulations with an integration time of 2 s. For Raman spectral images, spectra were collected at a step size of 1 µm in horizontal and vertical direction by an integration time of 0.25 s for each spectrum. Automated cosmic ray correction, background subtraction, and fitting using a Lorentz function were performed using the WITec Project software. Raman images were additionally processed by spectral averaging/smoothing and component analysis.

### 3.3 Stable isotope analyses (δ^13^C_carb_, δ^18^O_carb_)

Fifteen samples (ca. 100µg each) were extracted with a high-precision drill from individual mineral phases of polished rock slabs. The measurements were performed at 70 °C using a Thermo Scientific Kiel IV carbonate device coupled to a Finnigan DeltaPlus gas isotope mass spectrometer. Carbon and oxygen stable isotope ratios of carbonate minerals are reported as delta values (δ^13^C_carb_ and δ^18^O_carb_, respectively) relative to Vienna Pee Dee Belemnite (VPDB) reference standard. The standard deviation is 0.08‰ for δ^13^C_carb_ and 0.11‰ for δ^18^O_carb_.

All preparation and analytical work have been carried out at the Geoscience Center of the Georg-August-Universität Göttingen.

## 4 Results

### 4.1 Sedimentary succession in Jena and surroundings (Upper Buntsandstein, Olenekian, Early Triassic)

The Jena and surroundings section begins with a 1.5 m thick gypsum bed that contains no fossils (Bed 1) (Fig. 2). The gypsum bed is followed by a ∼3.5 m thick interval of grayish-green marls intercalated with two thin layers of dolomites (Bed 2) (Fig. 2). The marl interval is overlain by a 0.5 m thick bed of gray bioclastic limestone (*Tenuis*-bank) (Bed 3) which is marked by the first occurrence of the ammonoid *Beneckeia tenuis*. In addition to abundant ammonites, the *Tenuis*-bank contains various bivalves such as *Pseudomyoconcha gastrochaena, Hoernesia socialis, Neoschizodus elongatus, Neoschizodus ovatus* and *Costatoria costata*. It is directly followed by a 10 cm thick stromatolite bed (Fig. 2). This bed can be divided into a lower non-laminated and an upper laminated part (Fig. 2). The lamination is planar to wavy (Fig. 4a), but locally appears to be disrupted (Fig. 4b). The stromatolite bed consists mainly of dolomite as indicated by μ-XRF and Raman spectroscopy (Fig. 5). It is overlain by a ∼4.5 m thick interval of grayish-green sandy marl (Bed 4). The upper part of this interval contains a 0.6 m thick bed of grayish-green sandstone that contains fossils of various bivalves (*Pleuromya musculoides, Bakevellia mytiloides, C. costata*) and brachiopods (*Lingularia tenuissima*).

**Fig. 4.**
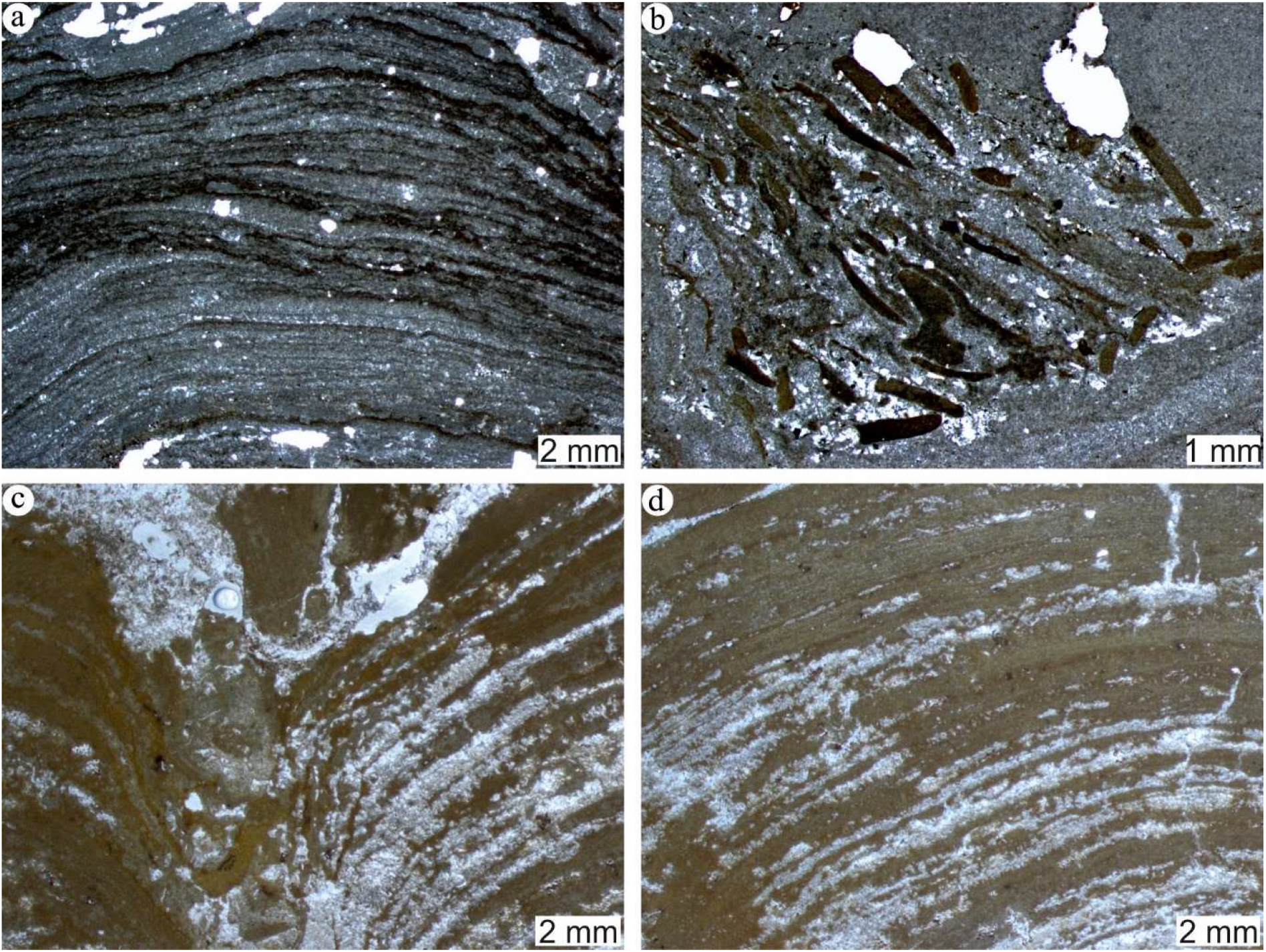
Thin section images (transmitted light) of stromatolites from Jena and surroundings (**a, b**) and Werbach (**c, d**). **a** Planar to wavy laminations in a Jena and surroundings stromatolite; **b** Disrupted laminations in a Jena and surroundings stromatolite; **c, d** Wavy to columnar laminations in a Werbach stromatolite

**Fig. 5.**
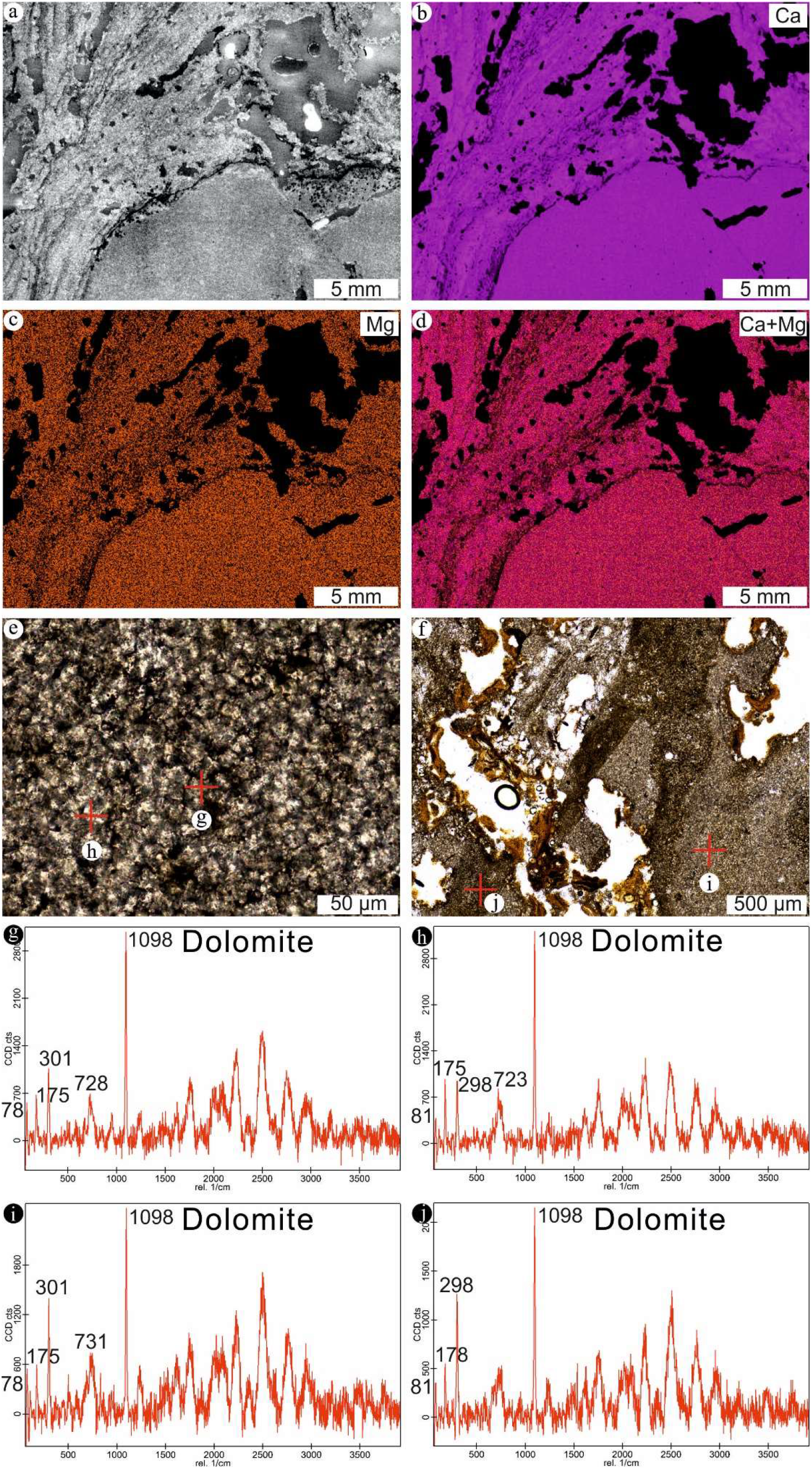
Micro X-ray fluorescence (μ-XRF) images and Raman spectroscopy data (single spectra) for a stromatolite from Jena and surroundings. **a** Scan image (reflected light); **b** Calcium (Ca) distribution; **c** Magnesium (Mg) distribution; **d** Calcium (Ca) plus Magnesium (Mg) distribution; **e, g, h** Raman spectra of dolomite in the lower non-laminated part of the stromatolite; **f, i, j** Raman spectra of dolomite in the upper laminated part of the stromatolite

The top half of the section starts with a ∼0.7 m thick bed of red platy sandstone (Bed 5), followed by a ∼1.3 m thick grayish-green marl bed (Bed 6) and a ∼0.1 m thick dolomite bed (Bed 7) (Fig. 2). The dolomite bed is overlain by a ∼0.9 m thick interval of bioclastic limestones (coquina) with abundant fossils (e.g., *C. costata, N. elongatus, B. tenuis*) and an oolite layer (Bed 8) (Fig. 2). After a thin marl layer, the succession continues with a ∼0.8 m thick red platy sandstone layer that shows wave ripple structures, desiccation cracks, and various types of bivalves (Bed 9) (Fig. 2). The section is terminated by a ∼6.5 m thick interval of grayish-green marl, intercalated with thin layers of sandstone, and gray dolomite (Bed 10) (Fig. 2). The dolomite layers exhibit wave ripples and contain (trace-) fossils (e.g., *C. costata, Rhizocorallium* sp.). The uppermost part of the grayish-green interval contains red gypsum nodules and (trace-) fossils (bivalve *Leptochondria albertii, C. costata, Rhizocorallium* sp.) (Fig. 2).

### 4.2 Sedimentary succession at Werbach (Middle Muschelkalk, Anisian, Middle Triassic)

In case of Werbach, our study focuses on the Karlstad Formation, which constitutes the lower part of the section. The relevant part begins with a ∼1.2 m thick bed of gray marl (Bed 1) (Fig. 3). This passes into a ∼1.2 m thick layer of ochre-colored dolomite and a ∼1m thick layer of ochre-colored dolomitic limestone with about 25 cm thick stromatolites (Bed 2) (Fig. 3). Bed 2 corresponds to the Geislingen Bed, a supraregional marker horizon (Simon et al. 2020). The stromatolites exhibit flat-columnar shapes (Fig. 3) and wavy to columnar laminations (Fig. 4c, d). They are mainly composed of calcite as revealed by μ-XRF and Raman spectroscopy (Fig. 6), but locally contain dolomite crystals (Fig. 6f, i). Following the dolomitic limestone with stromatolites, the section continues with a ∼6.5 m thick interval of alternating gray dolomite and marl layers (Bed 3). Bivalve fossils and intraclasts are observed at the base of this interval, whilst stromatolites can be found at the top (Fig. 3). The thickness of the above Heilbronn Formation is strongly reduced due to subsurface dissolution. It mainly consists of halite and gypsum, intercalated with dolomitic marls and limestones (Bed 4) (Fig. 3). The section ends with the Diemel Formation, characterized by dolomitic limestones (Bed 5) (Fig. 3).

**Fig. 6.**
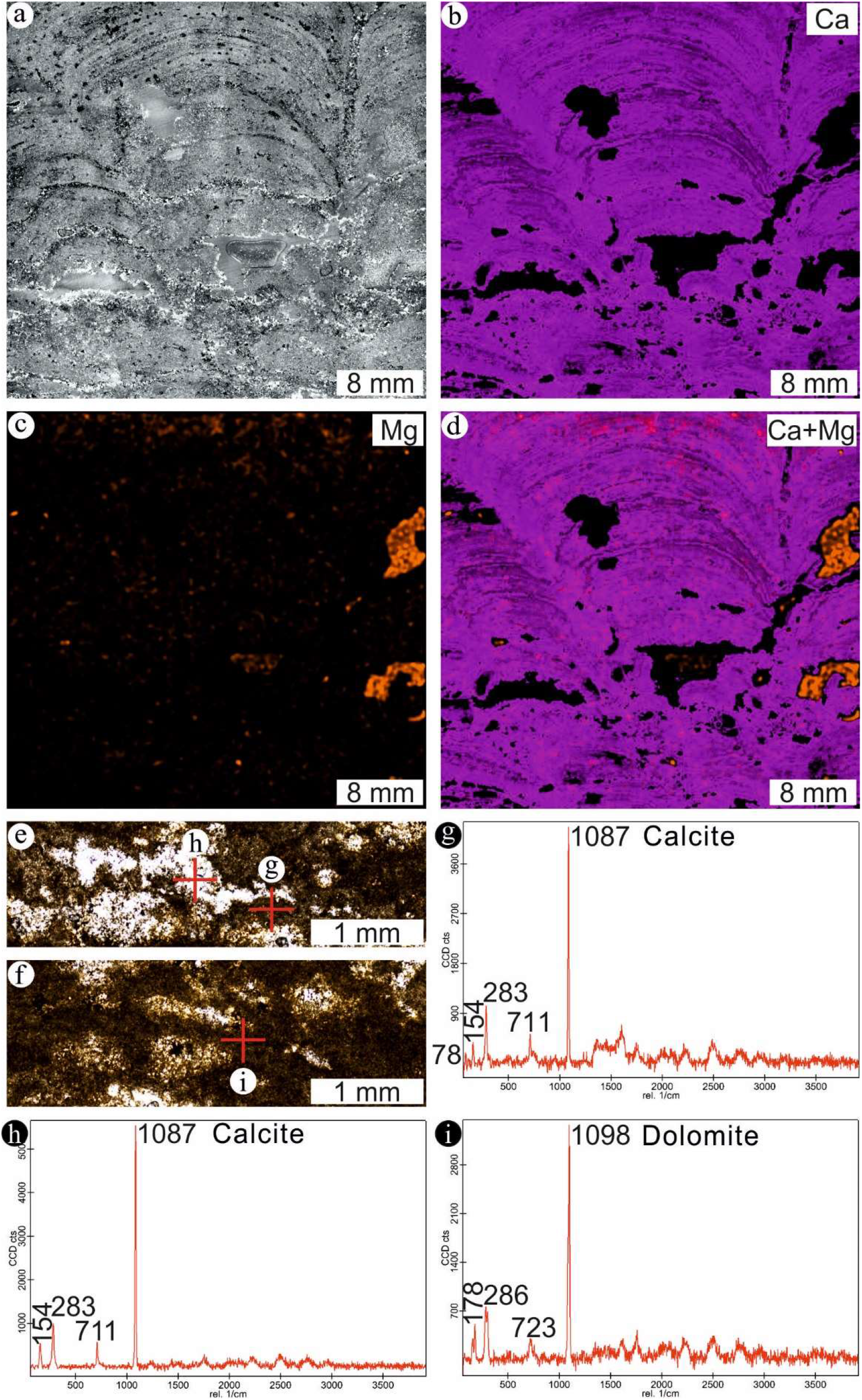
Micro X-ray fluorescence (μ-XRF) images and Raman spectroscopy data (single spectra) for a stromatolite from Werbach. **a** Scan image (reflected light); **b** Calcium (Ca) distribution; **c** Magnesium (Mg) distribution; **d** Calcium (Ca) plus Magnesium (Mg) distribution; **e, g, h** Raman spectra of calcite; **f, i** Raman spectra of dolomite

### 4.3 Hardheim microbe-metazoan build-up (Middle Muschelkalk, Anisian, Middle Triassic)

Regarding Hardheim, microbe-metazoan build-ups are ∼10 cm thick (Fig. 7a). The build-ups generally consist of calcite, dolomite, quartz, anatase and organic matter, as demonstrated by μ-XRF and Raman spectroscopy (Figs. 8, 9). Two groups of dolomite can be distinguished (Fig. 9c, d). The first group comprises euhedral dolomite crystals (Fig. 9c). The second group includes anhedral dolomite crystals that are intimately associated with organic matter (Fig. 9d). The microbe-metazoan build-ups are characterized by distinctly laminated columns (Fig. 7a). Laminations made of calcite with relatively abundant organic matter alternate with those made of calcite without organic matter (Fig. 8e, g, h). Non-spicular demosponges (“keratose”) can be found between and rarely within the columns (Fig. 7b-e). The sponges can clearly be distinguished by mesh-like fabrics and clotted to peloidal features (see Pei et al. 2021). In case of non-spicular demosponges, clotted to peloidal parts are composed of calcite and contain abundant organic matter (Fig. 8f, i), whilst areas characterized by mesh-like fabrics solely consist of calcite and are devoid of organic matter (Fig. 8f, j).

**Fig. 7.**
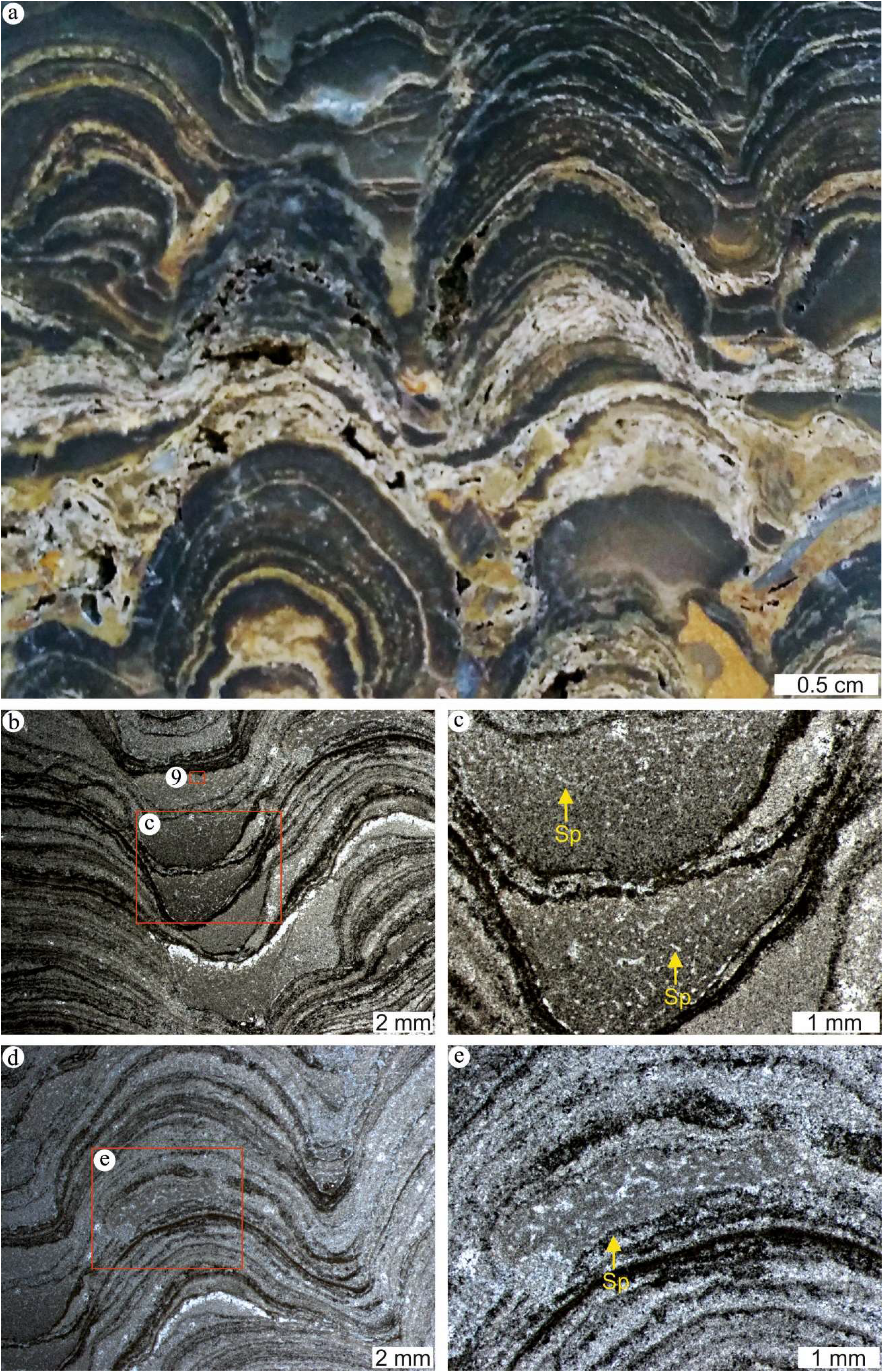
Overview photo and thin section images (transmitted light) of a microbe-metazoan build-up from Hardheim. **a** The build-up is mainly composed of columnar laminations; **b, c** Non-spicular demosponges between the laminated columns, showing characteristic mesh-like fabrics and clotted to peloidal features; **d, e** Non-spicular demosponges within laminated columns

**Fig. 8.**
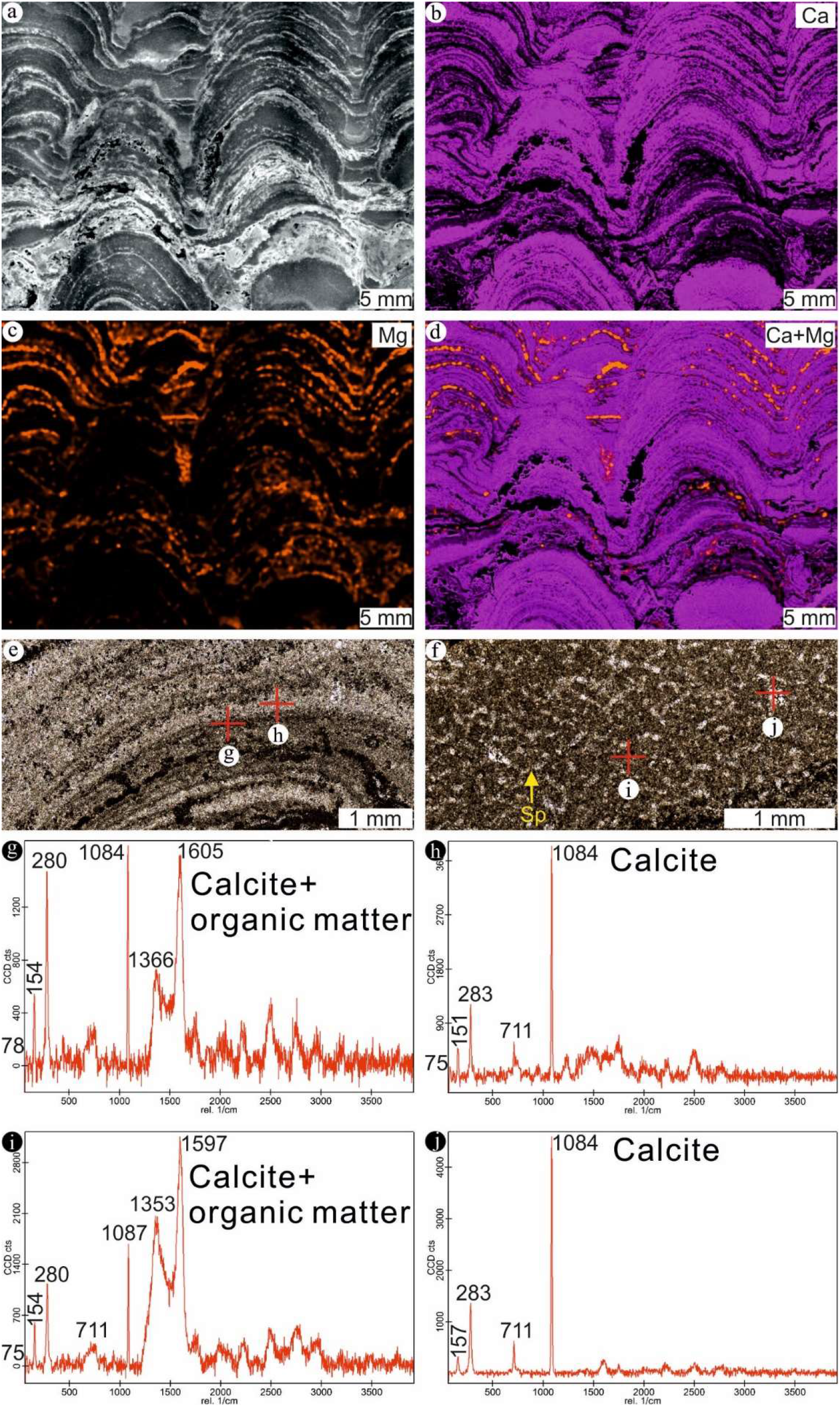
Micro X-ray fluorescence (μ-XRF) images and Raman spectroscopy data (single spectra) of a microbe-metazoan build-up from Hardheim. **a** Scan image (reflected light); **b** Calcium (Ca) distribution; **c** Magnesium (Mg) distribution; **d** Calcium (Ca) plus Magnesium (Mg) distribution; **e, g, h** Laminations made of calcite with relatively abundant organic matter alternate with those made of calcite without organic matter; **f, i, j** Clotted to peloidal parts of non-spicular demosponge fossils consist of calcite and contain abundant organic matter, while areas characterized by mesh-like fabrics solely consist of calcite and are devoid of organic matter

**Fig. 9.**
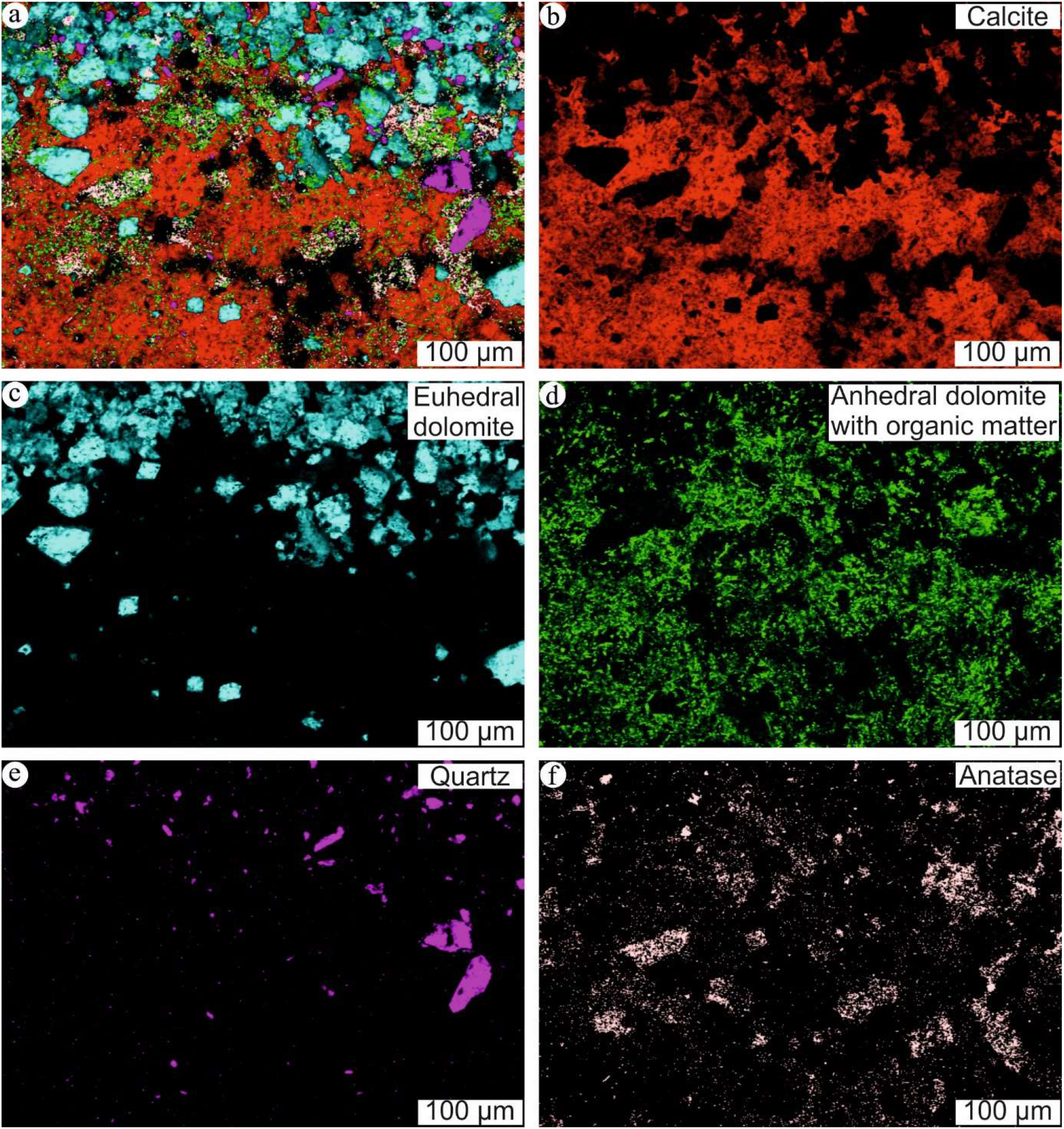
Raman spectroscopy data (spectral images) of a microbe-metazoan build-up from Hardheim (location indicated in 7**b**). **a** Combined image of calcite, dolomite, quartz, anatase and organic matter; **b** Distribution of calcite; **c** Distribution of euhedral dolomite; **d** Distribution of anhedral dolomite with organic matter; **e** Distribution of quartz; **f** Distribution of anatase

### 4.4 Carbon and oxygen stable isotopes (δ^13^C_carb_, δ^18^O_carb_)

δ^13^C_carb_ and δ^18^O_carb_ data for microbialites from the different localities cluster in three discrete groups (Fig. 10; Tab. 1). δ^13^C_carb_ and δ^18^O_carb_ values of Group 1 (Jena and surroundings stromatolite) range from −5.5 ‰ to −4.7 ‰ and −2.1 ‰ to −0.6 ‰, respectively. δ^13^C_carb_ values of Group 2 (Werbach stromatolite) vary between −5.8 ‰ and - 5.5 ‰, in the similar range with those of Group 1. δ^18^O_carb_ signatures of these samples, however, appear to be more negative, with an average value of −6.5 ‰. Marl samples from below the stromatolite layer at Werbach exhibits totally different δ^13^C_carb_ and δ^18^O_carb_ values, ranging from 1.3 ‰ to 1.5 ‰ and −2.0 ‰ to −1.9 ‰, respectively. δ^13^C_carb_ and δ^18^O_carb_ values of Group 3 (Hardheim microbe-metazoan build-up) vary from −1.5 ‰ to 0.6 ‰ and - 6.8 ‰ to −5.8 ‰, respectively. An extraclast contained in a microbe-metazoan build-up from Hardheim has a δ^13^C_carb_ value of −4.7 ‰ and a δ^18^O_carb_ value of −6.6 ‰ (Fig. 10; Tab. 1).

**Fig. 10.**
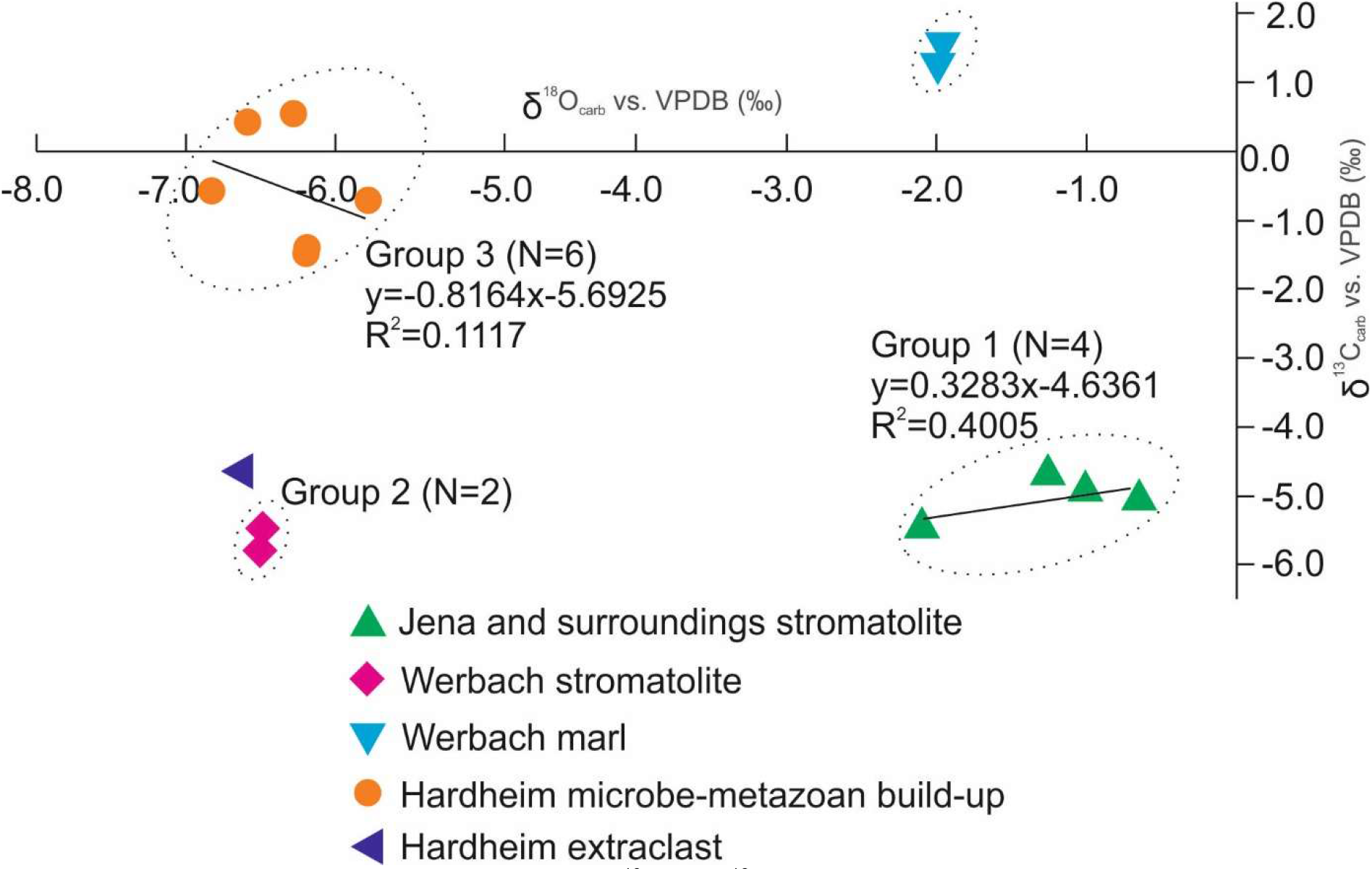
Carbon and oxygen stable isotope data (δ^13^C_carb_, δ^18^O_carb_, respectively) for microbialites from Jena and surroundings, Werbach, and Hardheim

**Tab. 1.** Carbon and oxygen stable isotope data (δ^13^C_carb_, δ^18^O_carb_, respectively) for microbialites from Jena and surroundings, Werbach and Hardheim

| Sample name | Sample number | $\delta^{13}\text{C}_{\text{carb}}$ vs. VPDB(‰) | $\delta^{18}\text{O}_{\text{carb}}$ vs. VPDB(‰) |
| --- | --- | --- | --- |
| Jena and surroundings stromatolite | 4 | -4.7 | -1.3 |
|  |  | -4.9 | -1.0 |
|  |  | -5.5 | -2.1 |
|  |  | -5.1 | -0.6 |
| Werbach stromatolite | 2 | -5.8 | -6.5 |
|  |  | -5.5 | -6.5 |
| Werbach marl | 2 | 1.3 | -2.0 |
|  |  | 1.5 | -1.9 |
| Hardheim microbe-metazoan build-up | 6 | -0.6 | -6.8 |
|  |  | -0.7 | -5.8 |
|  |  | -1.4 | -6.2 |
|  |  | -1.5 | -6.2 |
|  |  | 0.4 | -6.6 |
|  |  | 0.6 | -6.3 |
| Hardheim extraclast | 1 | -4.7 | -6.6 |

δ^13^C_carb_ and δ^18^O_carb_ values of Group 1 (Jena and surroundings stromatolite) show a relatively low coefficient of determination (R^2^ = 0.4005; N = 4) (Fig. 10), which might indicate a little effect of diagenetic alteration (Bishop et al. 2014). The R^2^ value for Group 3 (Hardheim microbe-metazoan build-up) is also low (0.1117; N = 6) and may be interpreted similarly (Fig. 10). However, the significantly negative δ^18^O_carb_ values of Group 3 perhaps reflect meteoric influence during diagenesis (Craig 1961), which could also be the case for Group 2 (Werbach stromatolite).

## 5 Discussion

### 5.1 Sedimentary environments

During Permian and Triassic times, the Germanic Basin was located on the edge of the western subtropical Tethys Ocean systems (Scotese and McKerrow 1990; Ziegler 1990; Stampfli 2000) (Fig. 1). In the Olenekian, a transgression from the western Tethys via the East Carpathian Gate resulted in the establishment of marine shelf environments in the surroundings of South Poland. Temporary, short-term transgressions entered the central Germanic basin, followed by dessication and playa deposits. In the area of Thuringia, this is reflected by the widespread deposition of marls, limestones, and dolomites, together with oolites and stromatolites. The sequence is formed by three cycles, each starting with marine transgressive deposition. The oolithic and bioclastic limestones are inferred to be formed in a barrier bar, open marine environment, leaving only thin veneers (lag deposits) behind. High sea-level is probably expressed by marls with an increasing content of sand to the top. Dessication cracks and gypsum nodules suggest a sabkha environment, before the next transgression started.

Strata of the Werbach section belong to the Middle Muschelkalk Subgroup (Anisian, Middle Triassic) and are thus stratigraphically younger than those exposed in Jena and surroundings (Fig. 3). Lithologically, the Werbach section comprises evaporites and carbonates such as dolomites, limestones, and marls (Simon et al. 2020) (Fig. 3). The lack of fossils except for local occurrences of hypersaline faunas (Simon et al. 2020) suggest hypersaline lagoon to sabkha environments. Since Hardheim is paleogeographically proximate to Werbach (Fig. 1), and the microbialites at both sections can be correlated stratigraphically (Simon et al. 2020), a similar paleoenvironment appears plausible. At the same time, different δ^13^C_carb_ values of stromatolites from Jena and surroundings, Werbach and microbe-metazoan build-ups from Hardheim might reflect differences in local evaporation, precipitation, and/or seawater influence, which is in good accordance with paleogeographic reconstructions (Ziegler 1990; Götz and Feist-Burkhardt 2012; Hagdorn 2020) (Fig. 1). Taken together, Lower-Middle Triassic microbialites probably formed in hypersaline lagoons to sabkha environments.

### 5.2 Stromatolites vs. microbe-metazoan build-ups

Olenekian stromatolites from Jena and surroundings exhibit planar to wavy laminations (Fig. 4a) and consist mainly of dolomite (Fig. 5). Anisian stromatolites from Werbach show wavy to columnar laminations (Fig. 4c, d) but are mainly composed of calcite (Fig. 6), which perhaps formed through dedolomitization (Hauck et al. 2018). Microbe-metazoan build-ups at Hardheim are characterized by columnar laminations (Fig. 7a) and consist mainly of calcite, dolomite, and organic matter (Figs. 8, 9). The distinct lamination textures result from the relative proportion of organic matter (Fig. 8e, g, h). Laminae with abundant organic matter likely represent exopolymeric substances (EPS) secreted by microbial mat communities (see Suarez-Gonzalez and Reitner 2021). Those devoid of organic matter, in contrast, might be dominated by authigenic mineral precipitation and/or trapping and binding of detrital materials (Awramik et al. 1976; Reitner 2011; Suarez-Gonzalez et al. 2019).

The major difference between all the studied microbialites is the presence of non-spicular demosponges in the microbe-metazoan build-ups from Hardheim. As discussed above, non-spicular demosponges occur between and within laminated columns, and are readily discernable by mesh-like fabrics and clotted to peloidal features (Fig. 7b-e). Parts characterized by mesh-like fabrics solely consist of calcite (Fig. 8f, j) while those exhibiting clotted to peloidal features also contain organic matter (Fig. 8f, i). Three-dimensional reconstructions of modern and fossil non-spicular demosponges establish that the mesh-like fabrics represent skeletal elements that originally consist of spongin/chitin (Luo and Reitner 2014, 2016; see Pei et al. 2021, their Fig. 1). The clotted to peloidal features, in contrast, reflect automicrite that form through the *in-situ* microbial decay of microbe-rich sponge tissue (Reitner 1993; Reitner et al. 1995).

Notably, anhedral dolomite crystals in microbial-metazoan build-ups from Hardheim are intimately associated with organic matter (Fig. 9d). They are tentatively attributed to protodolomite, hypothetically related to microbial sulfate reduction (Warthmann et al. 2000; Liu et al. 2020). In particular, various modern demosponges (e.g., *Chondrosia reniformis, Petrosia ficiformis and Geodia barretti*) harbor abundant sulfate reducing bacteria, and pyrite crystals, the typical end product of microbial sulfate reduction, are commonly present in taphonomically mineralized sponge tissue (Reitner and Schumann-Kindel 1997; Schumann-Kindel et al. 1997; Hoffmann et al. 2005; Zhang et al. 2015). It is thus tempting to speculate that the anhedral dolomite crystals in microbe-metazoan build-ups from Hardheim resulted from taphonomic processes associated with the sponges.

### 5.3 Paleoecological implications of the microbialites

The Permian – Triassic crisis is distinguished by a potentially catastrophic decline of biodiversity in marine and terrestrial ecosystems (Benton and Twitchett 2003; Wignall 2007; Chen and Benton 2012; Payne and Clapham 2012). Furthermore, it is associated with ubiquitous occurrences of unusual sedimentary features, including microbialites (Woods 2014; Wu et al. 2017; Heindel et al. 2018; Chen et al. 2019; Pei et al. 2019). As stated before, microbe-metazoan build-ups are easily overlooked in geological records, but have received much attention and interests recently (Luo and Reitner 2014, 2016; Friesenbichler et al. 2018; Heindel et al. 2018; Baud et al. 2021; Lee and Riding 2021; Pei et al. 2021).

One possible explanation for the widespread occurrence of diverse microbial mats is a suppressed ecological competition with grazing metazoans that would prevent their development (Foster et al. 2020). Although the details remain to be studied, it seems plausible that microbes and non-spicular demosponges had a mutualistic relationship (Luo and Reitner 2016; Lee and Riding 2021; Pei et al. 2021), and it is tempting to speculate that the investigated microbial-metazoan build-ups reflect an ancient evolutionary and ecologic association.

Salinity appears to be an additional influential factor. Microbial mats can develop under salinities of up to 170 ‰ (e.g., Schneider et al. 2013), but most animals suffer from high salinities (Bayly 1972). Notably, various sponges are more tolerant (e.g., *Microciona prolifera*), capable of surviving in environments with salinities up to 45 ‰ (e.g., Leamon and Fell 1990). The Lower-Middle Triassic microbialites studied herein formed in hypersaline lagoon to sabkha environments. Perhaps it was minor differences in salinities that controlled whether or not non-spicular demosponges could develop. This may explain the preferential development of non-spicular sponges in morphological valleys between laminated columns, since these areas might have been characterized by slightly lower salinities as compared to the top parts of the columns.

## 6 Conclusion

Triassic microbialites from Jena and surroundings (Upper Buntsandstein Subgroup, Olenekian, Early Triassic) as well as from Werbach and Hardheim (both lower Middle Muschelkalk Subgroup, Anisian, Middle Triassic) formed in hypersaline lagoons to sabkha environments. Olenekian stromatolites in Jena and surroundings exhibit planar to wavy laminations. Anisian stromatolites from Werbach are characterized by wavy to columnar laminations. Anisian microbialites from Hardheim (microbe-metazoan build-ups) consist of columnar laminations, which are different in that they embrace non-spicular demosponges. Stromatolite formation relies on authigenic mineral precipitation controlled by exopolymeric substances secreted by microbes in the mats and/or trapping and binding of detrital sediments. Skeletons of non-spicular demosponges originally consist of spongin/chitin. These organisms preserve through organomineralization processes linked to the microbial degradation of sponge tissue. This specific taphonomic process results in the formation of clotted to peloidal features that are characteristic for fossil non-spicular demosponges. The proliferation of microbial mats and/or microbe-metazoan build-ups was likely due to the suppressed ecological competition after the Permian – Triassic crisis. It appears plausible that microbes and non-spicular demosponges in the build-ups had a mutualistic relationship, and it is tempting to speculate that this association reflects an ancient evolutionary and ecologic strategy. Given the paleoenvironments, salinity might have been another ecological control on the presence of non-spicular demosponges.

## Abbreviations

μ-XRF: Micro X-ray fluorescence

## 7 Supplementary Information

## Acknowledgements

Axel Hackmann, Burkhard Schmidt, Dennis Kohl, Jan Schönig and Thierry Wasselin are thanked for lab assistance. Thomas Billert is thanked for providing stromatolites from Jena and surroundings.

## Authors’ contributions

YP and JR designed the study. HH and TV provided the sections and samples. YP, JR, HH, TV and JPD carried out discussions throughout the whole process. YP conducted sample analyses and wrote the original draft. YP, JPD, JR, HH and TV revised the draft. The final draft was approved by YP, JR, JPD, HH and TV.

## Founding

This study was financially supported by the China Scholarship Council.

## Availability of data and materials

All research data and results are displayed in this published article.

## References

Aitken, J.D. 1966. Middle Cambrian to Middle Ordovician cyclic sedimentation, southern Rocky Mountains of Alberta. Bulletin of Canadian Petroleum Geology 14: 405–441. https://doi.org/10.35767/gscpgbull.14.4.405.

Arp, G., A. Reimer, and J. Reitner. 2001. Photosynthesis-induced biofilm calcification and calcium concentrations in Phanerozoic oceans. Science 292 (5522): 1701–1704. doi: 10.1126/science.1057204.

Arp, G., C. Ostertag-Henning, S. Yücekent, J. Reitner, and V. Thiel. 2008. Methane-related microbial gypsum calcitization in stromatolites of a marine evaporative setting (Münder Formation, Upper Jurassic, Hils Syncline, north Germany). Sedimentology 55 (5): 1227–1251. https://doi.org/10.1111/j.1365-3091.2007.00944.x.

Arp, G., V. Thiel, A. Reimer, W. Michaelis, and J. Reitner. 1999. Biofilm exopolymers control microbialite formation at thermal springs discharging into the alkaline Pyramid Lake, Nevada, USA. Sedimentary Geology 126 (1–4): 159–176. https://doi.org/10.1016/S0037-0738(99)00038-X.

Awramik, S.M., L. Margulis, and E.S. Barghoorn. 1976. Evolutionary processes in the formation of stromatolites. In Stromatolites, ed. M.R. Walter. Developments in Sedimentology 20: 149–162. Amsterdam: Elsevier. https://doi.org/10.1016/S0070-4571(08)71135-X.

Bachmann, G.H. 2002. A Lamellibranch-stromatolite bioherm in the Lower Keuper (Ladinian, Middle Triassic), South Germany. Facies 46: 83–88.

Bachmann, G.H. and Gwinner, M.P. 1971. Algen-Stromatolithen von der Grenze Unterer/Mittlerer Keuper (Obere Trias) bei Schwäbisch Hall (Nordwürttemberg, Deutschland). Neues Jahrbuch für Geologie und Paläontologie, Monatshefte 1971: 594–604.

Bayly, I.A.E. 1972. Salinity tolerance and osmotic behavior of animals in athalassic saline and marine hypersaline waters. Annual review of ecology and systematics 3: 233–268.

Baud, A., S. Richoz, R. Brandner, L. Krystyn, K. Heindel, T. Mohtat, P. Mohtat-Aghai, and M. Horacek. 2021. Sponge takeover from end-Permian mass extinction to Early Induan time: Records in Central Iran microbial buildups. Frontiers in Earth Science 9: 586210. doi: 10.3389/feart.2021.586210.

Benton, M.J., and R.J. Twitchett. 2003. How to kill (almost) all life: the end-Permian extinction event. Trends in Ecology and Evolution 18 (7): 358–365. https://doi.org/10.1016/S0169-5347(03)00093-4.

Bishop, J.W., D.A. Osleger, I.P. Montañez, and D.Y. Sumner. 2014. Meteoric diagenesis and fluid-rock interaction in the Middle Permian Capitan backreef: Yates Formation, Slaughter Canyon, New Mexico. American Association of Petroleum Geologists, Bulletin 98 (8): 1495–1519. https://doi.org/10.1306/05201311158.

Bosak, T., B. Liang, T.-D. Wu, S.P. Templer, A. Evans, H. Vali, J.-L. Guerquin-Kern, V. Klepac-Ceraj, M.S. Sim, and J. Mui. 2012. Cyanobacterial diversity and activity in modern conical microbialites. Geobiology 10: 384– 401. doi: 10.1111/j.1472-4669.2012.00334.x

Böhm, F., and T.C. Brachert. 1993. Deep-water stromatolites and Frutexites Maslov from the Early and Middle Jurassic of S-Germany and Austria. Facies 28: 145–168.

Braga, J.C., J.M. Martin, and R. Riding. 1995. Controls on microbial dome fabric development along a carbonate-siliciclastic shelf-basin transect, Miocene, SE Spain. Palaios 10: 347–361. https://doi.org/10.2307/3515160.

Burne, R.V., and L.S. Moore. 1987. Microbialites; organosedimentrary deposits of benthic microbial communities. Palaios 2 (3): 241–254. https://doi.org/10.2307/3514674.

Craig, H. 1961. Isotopic variations in meteoric waters. Science 133 (3465): 1702–1703. doi: 10.1126/science.133.3465.1702.

Chen, Z-Q., C. Tu, Y. Pei, J. Ogg, Y. Fang, S. Wu, X. Feng, Y. Huang, Z. Guo, and H. Yang. 2019. Biosedimentological features of major microbe-metazoan transitions (MMTs) from Precambrian to Cenozoic. Earth-Science Reviews 189: 21–50. https://doi.org/10.1016/j.earscirev.2019.01.015.

Chen, Z-Q., and M.J. Benton. 2012. The timing and pattern of biotic recovery following the end-Permian mass extinction. Nature Geoscience 5: 375–383. doi: 10.1038/NGEO1475.

Dravis, J.J. 1983. Hardened subtidal stromatolites, Bahamas. Science 219 (4583): 385–386. doi: 10.1126/science.219.4583.385.

Duda, J-P., M.J. Van Kranendonk, V. Thiel, D. Ionescu, H. Strauss, N. Schäfer, and J. Reitner. 2016. A Rare glimpse of Paleoarchean life: Geobiology of an exceptionally preserved microbial mat facies from the 3.4 Ga Strelley Pool Formation, Western Australia. Plos One 11 (1): e0147629. https://doi.org/10.1371/journal.pone.0147629.

Foster, W.J., K. Heindel, S. Richoz, J. Gliwa, D.J. Lehrmann, Baud. A, T. Kolar-Jurkovšek, D. Aljinović, B. Jurkovšek, D. Korn, R.C. Martindale, and J. Peckmann. 2020. Suppressed competitive exclusion enabled the proliferation of Permian/Triassic boundary microbialites. The Depositional Record 6: 62–74. https://doi.org/10.1002/dep2.97.

Friesenbichler, E., S. Richoz, A. Baud, L. Krystyn, L. Sahakyan, S. Vardanyan, J. Peckmann, J. Reitner, and K. Heindel. 2018. Sponge-microbial build-ups from the lowermost Triassic Chanakhchi section in southern Armenia: Microfacies and stable carbon isotopes. Palaeogeography, Palaeoclimatology, Palaeoecology 490: 653–672. https://doi.org/10.1016/j.palaeo.2017.11.056.

Genser, C. 1930. Zur Stratigraphie und Chemie des Mittleren Muschelkalks in Franken. Geologisch und paläontologische Abhandlungen, Neue Folge 17: 111 pp.

Götz, A.E., and S. Feist-Burkhardt. 2012. Phytoplankton associations of the Anisian Peri-Tethys Basin (Central Europe): Evidence of basin evolution and palaeoenvironmental change. Palaeogeography, Palaeoclimatology, Palaeoecology 337–338: 151–158. https://doi.org/10.1016/j.palaeo.2012.04.009.

Hagdorn, H. 2020. Paläobiogeographie des Mitteleuropäischen Beckens in der Frühen und Mittleren Trias und Faunenimmigration ins Muschelkalkmeer. In Stratigraphie von Deutschland XIII. Muschelkalk, ed. Deutsche Stratigraphische Kommission (Koordination und Redaktion: H. Hagdorn, and T. Simon für die Subkommission Perm-Trias). Schriftenreihe Deutsche Gesellschaft für Geowissenschaften 91: 111–123.

Hagdorn, H., and T. Simon. 1993. Rinnenbildung und Emersion in den Basisschichten des Mittleren Muschelkalks von Eberstadt (Nordbaden). Neues Jahrbuch für Geologie und Paläontologie 189: 119–145.

Hauck, T.E., H.J. Corlett, M. Grobe, E.L. Walton, and P. Sansjofre. 2018. Meteoric diagenesis and dedolomite fabrics in precursor primary dolomicrite in a mixed carbonate–evaporite system. Sedimentology 65: 1827–1858. doi: 10.1111/sed.12448.

Heim, C., K. Simon, D. Ionescu, A. Reimer, D. De Beer, N-V. Quéric, J. Reitner, and V. Thiel. 2015. Assessing the utility of trace and rare earth elements as biosignatures in microbial iron oxyhydroxides. Frontiers in Earth Science 3: 1–15. https://doi.org/10.3389/feart.2015.00006.

Heim, C., N.-V. Quéric, D. Ionescu, N. Schäfer, and J. Reitner. 2017. Frutexites-like structures formed by iron oxidizing biofilms in the continental subsurface (Äspö Hard Rock Laboratory, Sweden). PLoS ONE 12 (5): e0177542. https://doi.org/10.1371/journal.pone.0177542.

Heindel, K., W.J. Foster, S. Richoz, D. Birgel, V.J. Roden, A. Baud, R. Brandner, L. Krystyn, T. Mohtat, E. Koşun, R.J. Twitchett, J. Reitner, and J. Peckmann. 2018. The formation of microbial-metazoan bioherms and biostromes following the latest Permian mass extinction. Gondwana Research 61: 187–202. https://doi.org/10.1016/j.gr.2018.05.007.

Hoffmann, F., O. Larsen, V. Thiel, H.T. Rapp, T. Pape, W. Michaelis, and J. Reitner. 2005. An anaerobic world in sponges. Geomicrobiology Journal 22 (1–2): 1–10. https://doi.org/10.1080/01490450590922505.

Kalkowsky, E. 1908. Oolith und Stromatolith im norddeutschen Buntsandstein. Zeitschrift der Deutschen Geologischen Gesellschaft 60: 68–125.

Leamon, J., and P.E. Fell. 1990. Upper salinity tolerance of and salinity-induced tissue regression in the estuarine sponge Microciona prolifera. Transactions of the American Microscopical Society 109 (3): 265–272. https://www.jstor.org/stable/3226797.

Lee, J-H., and R. Riding. 2021. The ‘classic stromatolite’ Cryptozoön is a keratose sponge-microbial consortium. Geobiology 19: 189–198. https://doi.org/10.1111/gbi.12422.

Liu, D., Q. Fan, D. Papineau, N. Yu, Y. Chu, H. Wang, X. Qiu, and X. Wang. 2020. Precipitation of protodolomite facilitated by sulfate-reducing bacteria: The role of capsule extracellular polymeric substances. Chemical Geology 533: 119415. https://doi.org/10.1016/j.chemgeo.2019.119415.

Luo, C., and J. Reitner. 2014. First report of fossil “keratose” demosponges in Phanerozoic carbonates: preservation and 3-D reconstruction. Naturwissenschaften 101: 467–477. doi: 10.1007/s00114-014-1176-0.

Luo, C., and J. Reitner. 2016. ‘Stromatolites’ built by sponges and microbes – a new type of Phanerozoic bioconstruction. Lethaia 49: 555–570. https://doi.org/10.1111/let.12166.

Mißbach, H., J-P. Duda, A.M. van den Kerkhof, V. Lüders, A. Pack, J. Reitner, and V. Thiel. 2021. Ingredients for microbial life preserved in 3.5 billion-year-old fluid inclusions. Nature Communications 12: 1–11. https://doi.org/10.1038/s41467-021-21323-z.

Naumann, E. 1928. Erläuterungen zur Geologischen Karte von Preußen und benachbarten deutschen Ländern. Blatt Jena, Gradabteilung 71, Blatt 2, - 65 S, Berlin.

Paul, J., T.M. Peryt, and R.V. Burne. 2011. Kalkowsky’s stromatolites and oolites (Lower Buntsandstein, Northern Germany). In Advances in Stromatolite Geobiology, ed. J. Reitner, N.-V. Quéric, and G. Arp: 13–28. Berlin: Springer.

Payne, J.L., and M.E. Clapham. 2012. End-Permian Mass Extinction in the Oceans: An Ancient Analog for the Twenty-First Century? Annual Review of Earth and Planetary Sciences 40: 89–111. https://doi.org/10.1146/annurev-earth-042711-105329.

Pei, Y., J.-P. Duda, J. Schönig, C. Luo, and J. Reitner. 2021. Late Anisian microbe-metazoan build-ups (‘stromatolites’) in the Germanic Basin — aftermath of the Permian — Triassic Crisis. doi: 10.1101/2021.03.15.435468.

Pei, Y., Z-Q. Chen, Y. Fang, S. Kershaw, S. Wu, and M. Luo. 2019. Volcanism, redox conditions, and microbialite growth linked with the end-Permian mass extinction: Evidence from the Xiajiacao section (western Hubei Province), South China. Palaeogeography, Palaeoclimatology, Palaeoecology 519: 194–208. https://doi.org/10.1016/j.palaeo.2017.07.020.

Playford, P.E., A.E. Cockbain, E.C. Druce, and J.L. Wray. 1976. Devonian stromatolites from the Canning Basin, Western Australia. doi: In Stromatolites, ed. M.R. Walter. Developments in Sedimentology 20: 543–563. Amsterdam: Elsevier. https://doi.org/10.1016/S0070-4571(08)71158-0.

Reitner, J. 1993. Modern cryptic microbialite/metazoan facies from Lizard Island (Great Barrier Reef, Australia) formation and concepts. Facies 29: 3–39.

Reitner, J. 2011. Microbial mats. In Encyclopedia of Geobiology, ed. J. Reitner, and V. Thiel: 606–608. Berlin: Springer.

Reitner, J., and G. Schumann-Kindel. 1997. Pyrite in mineralized sponge tissue – product of sulfate reducing sponge related bacteria? In Biosedimentology of microbial buildups IGCP Project No. 380, ed. F. Neuweiler, J. Reitner, and C. Monty. Facies 36: 272–284.

Reitner, J., J. Peckmann, A. Reimer, G. Schumann, and V. Thiel. 2005b. Methane-derived carbonate build-ups and associated microbial communities at cold seeps on the lower Crimean shelf (Black Sea). Facies 51: 66–79. doi 10.1007/s10347-005-0059-4.

Reitner, J., J. Peckmann, M. Blumenberg, W. Michaelis, A. Reimer, and V. Thiel. 2005a. Concretionary methane-seep carbonates and associated microbial communities in Black Sea sediments. Palaeogeography, Palaeoclimatology, Palaeoecology 227 (1–3): 18–30. https://doi.org/10.1016/j.palaeo.2005.04.033.

Reitner, J., P. Gautret, F. Marin, and F. Neuweiler. 1995. Automicrite in a modern marine microbialite. Formation model via organic matrices (Lizard Island, Great Barrier Reef, Australia). Bulletin de l’Institut Océanographique (Monaco) Numéro Spécial 14: 237–263.

Riding, R. 1991. Classification of microbial carbonates. In Calcareous algae and stromatolites, ed. R. Riding: 21–51. Berlin: Springer.

Riding, R. 2006. Microbial carbonate abundance compared with fluctuations in metazoan diversity over geological time. Sedimentary Geology 185 (3–4): 229–238. https://doi.org/10.1016/j.sedgeo.2005.12.015.

Riding, R., and L. Liang. 2005. Geobiology of microbial carbonates: metazoan and seawater saturation state influences on secular trends during the Phanerozoic. Palaeogeography, Palaeoclimatology, Palaeoecology 219 (1–2): 101–115. https://doi.org/10.1016/j.palaeo.2004.11.018.

Stampfli, G.M. 2000. Tethyan oceans. In Tectonics and Magmatism in Turkey and the Surrounding Area, ed. E. Bozkurt, J.A. Winchester, and J.D.A. Piper. Geological Society, London, Special Publications 173: 1–23. https://doi.org/10.1144/GSL.SP.2000.173.

Szulc, J. 1997. Middle Triassic (Muschelkalk) sponge-microbial stromatolites, diplopores and Girvanella-oncoids from the Silesian Cracow Upland. In 3rd IFAA Regional Symposium and IGCP 380 International Meeting, Guidebook: 10–15. Cracow.

Schneider, D., G. Arp, A. Reimer, J. Reitner, and R. Daniel. 2013. Phylogenetic analysis of a microbialite-forming microbial mat from a hypersaline lake of the Kiritimati Atoll, Central Pacific. PLoS ONE 8 (6): e66662. https://doi.org/10.1371/journal.pone.0066662.

Schumann-Kindel, G., M. Bergbauer, W. Manz, U. Szewzyk, and J. Reitner. 1997. Aerobic and anaerobic microorganisms in modern sponges: a possible relationship to fossilization-processes. In Biosedimentology of microbial buildups IGCP Project No. 380, ed. F. Neuweiler, J. Reitner, and C. Monty. Facies 36: 268–272.

Schwarz, H.-U. 1970. Zur Sedimentologie und Fazies des Unteren Muschelkalkes in Südwestdeutschland und angrenzenden Gebieten. Unpublished Ph.D. Thesis. Eberhard-Karls-Universität Tübingen, Tübingen: 297 pp.

Scotese, C.R., and W.S. McKerrow. 1990. Revised world maps and introduction. Geological Society, London, Memoirs 12: 1–21. https://doi.org/10.1144/GSL.MEM.1990.012.01.01.

Simon, T., H. Hagdorn, D. Dittrich, S. Röhling, and T. Voigt. 2020. Lithostratigraphie der Mittlerer-Muschelkalk-Subgruppe. Schriftenreihe der Deutschen Gesellschaft für Geowissenschaften 91: 441–466.

Suarez-Gonzalez, P., and J. Reitner. 2021. Ooids forming in situ within microbial mats (Kiritimati atoll, central Pacific). doi: https://doi.org/10.1101/2021.05.05.442839.

Suarez-Gonzalez, P., M.I. Benito, I.E. Quijada, R. Mas, and S. Campos-Soto. 2019. ‘Trapping and binding’: A review of the factors controlling the development of fossil agglutinated microbialites and their distribution in space and time. Earth-Science Reviews 194: 182–215. https://doi.org/10.1016/j.earscirev.2019.05.007.

Vossmerbäumer, H. 1971. Organosedimentäre Gefüge im oberen Wellenkalk (mu3, Trias) Frankens. Neues Jahrbuch für Geologie und Paläontologie, Monatshefte 1971: 437–448.

Walter, M.R. 1972. Stromatolites and the biostratigraphy of the Australian Precambrian and Cambrian. Special Papers in Palaeontology 11: 1–256.

Woods, A.D. 2014. Assessing Early Triassic paleoceanographic conditions via unusual sedimentary fabrics and features. Earth-Science Reviews 137: 6–18. https://doi.org/10.1016/j.earscirev.2013.08.015.

Warthmann, R., Y. Van Lith, C. Vasconcelos, J.A. McKenzie, and A.M. Karpoff. 2000. Bacterially induced dolomite precipitation in anoxic culture experiments. Geology 28 (12): 1091–1094. https://doi.org/10.1130/0091-7613(2000)28<1091:BIDPIA>2.0.CO;2.

Wignall, P.B. 2007. The End-Permian mass extinction – how bad did it get? Geobiology 5: 303–309. https://doi.org/10.1111/j.1472-4669.2007.00130.x.

Wu, S., Z-Q. Chen, Y. Fang, Y. Pei, and H. Yang. 2017. A Permian–Triassic boundary microbialite deposit from the eastern Yangtze Platform (Jiangxi Province, South China): Geobiologic features, ecosystem composition, and redox conditions. Palaeogeography, Palaeoclimatology, Palaeoecology 486: 58–73. https://doi.org/10.1016/j.palaeo.2017.05.015.

Ziegler, P.A. 1990. Geological atlas of Western and Central Europe: 1–239. Bath: Geological Society Publishing House.

Zhang, D., W. Sun, G. Feng, F. Zhang, R. Anbuchezhian, Z. Li, and Q. Jiang. 2015. Phylogenetic diversity of sulphate-reducing Desulfovibrio associated with three South China Sea sponges. Letters in Applied Microbiology 60: 504–512. doi:10.1111/lam.12400.

